# Genetic Risk Scores for Cardiometabolic Traits in Sub-Saharan African Populations

**DOI:** 10.1101/2020.05.21.109199

**Authors:** Kenneth Ekoru, Adebowale A. Adeyemo, Guanjie Chen, Ayo P. Doumatey, Jie Zhou, Amy R. Bentley, Daniel Shriner, Charles N. Rotimi

## Abstract

There is growing support for the use of genetic risk scores (GRS) in routine clinical settings. Due to the limited diversity of current genomic discovery samples, there are concerns that the predictive power of GRS will be limited in non-European ancestry populations. Here, we evaluated the predictive utility of GRS for 12 cardiometabolic traits in sub-Saharan Africans (AF; *n*=5200), African Americans (AA; *n*=9139), and European Americans (EA; *n*=9594). GRS were constructed as weighted sums of the number of risk alleles. Predictive utility was assessed using the additional phenotypic variance explained and increase in discriminatory ability over traditional risk factors (age, sex and BMI), with adjustment for ancestry-derived principal components. Across all traits, GRS showed upto a 5-fold and 20-fold greater predictive utility in EA relative to AA and AF, respectively. Predictive utility was most consistent for lipid traits, with percent increase in explained variation attributable to GRS ranging from 10.6% to 127.1% among EA, 26.6% to 65.8% among AA, and 2.4% to 37.5% among AF. These differences were recapitulated in the discriminatory power, whereby the predictive utility of GRS was 4-fold greater in EA relative to AA and up to 44-fold greater in EA relative to AF. Obesity and blood pressure traits showed a similar pattern of greater predictive utility among EA. This work demonstrates the poorer performance of GRS in AF and highlights the need to improve representation of multiethnic populations in genomic studies to ensure equitable clinical translation of GRS.

**Key Messages:**

- Genetic Risk Score (GRS) prediction is markedly poorer in sub-Saharan Africans compared with African Americans and European Americans
- To ensure equitable clinical translation of GRS, there is need need to improve representation of multiethnic populations in genomic studies

## Background

The use of aggregate genetic risk, as summed up in genetic risk scores (GRS), to identify subgroups of individuals at increased risk of disease or more likely to benefit from early intervention, is gaining recognition as a practical translational strategy of genomic findings for both public health and clinical care. This trend is supported by evidence showing that risk associated with GRS for certain common complex diseases, such as severe obesity and coronary artery disease, can be as high as the risk conferred by some rare monogenic mutations, and that incorporating such GRS in disease risk prediction models can substantially increase prediction accuracy.^1–4^ However, GRS derived from existing genome-wide association studies (GWAS) show greater predictive value in European populations than in non-European populations, a reflection of the fact that most GWAS have been conducted in European-ancestry populations. For example, GRS derived from the largest available datasets show up to 2- to 5-fold greater predictive power in European-ancestry populations relative to African Americans and East Asians for a number of complex traits including anthropometric indices and mental health disorders.^5–8^

There are concerns that the adoption of routine use of GRS in clinical setting could exercebate existing health diaprities because of suboptimal utility in non-European-ancestry populations. Therefore, as the use of GRS moves from research to clinical settings, it is essential to clarify its utility in populations that are currently underrepresented in genomic discoveries. While there are limited data on the predictive utility of GRS in populations such as East Asians and African Americans, similar information is lacking in populations from continental Africa.^6–9^ In the present study, we sought to assess the predictive utility of GRS for a range of cardiometabolic traits in sub-Saharan Africans (AF) and make comparisons with European Americans (EA) and African Americans (AA). We aimed to do this using GRS constructed from genetic variants reported in publically available databases of GWAS to exemplify the potential use of such resources.

## Methods

### Study participants

The predictive utility of GRS was assessed in up to 5200 sub-Saharan Africans (AF), 9139 African Americans (AA), and 9594 individuals of European Americans (EA). AF were drawn from the AADM study ^10,11^ that enrolled participants aged 18 years or older from Nigeria, Ghana and Kenya as described previously .^12^ Data on AA were obtained from the Howard University Family Study (HUFS)^13^, and from the following dbGAP studies: Cleveland Family Study (CFS, phs000284)^14^, Jackson Heart Study (JHS, phs000286)^15^, Multi-Ethnic Study of Atherosclerosis (MESA, phs000209)^16^ and Atherosclerosis Risk in Communities Cohort (ARIC, phs000280)^17^;. CFS, JHS, HUFS, MESA and ARIC participants are aged 35-84 years and were recruited from different parts of the United States. Data on EA were obtained from the ARIC study.^17^

### Cardiometabolic traits studied

We studied body mass index (BMI), waist circumference (WC), hip circumference (HC), waist-to-hip ratio (WHR), systolic blood pressure (SBP), diastolic blood pressure (DBP), fasting plasma glucose (FPG), triglycerides (TG), total cholesterol (TC), low-density lipoprotein (LDL) and high-density lipoprotein (HDL), all measured in standard units; type 2 diabetes (T2D) status was determined according to the American Diabetes Association criteria. Additionally, we derived the following binary traits based on commonly used clinical definitions: general obesity (BMI ≥ 30 Kg/m^2^), abdominal obesity (WC: ≥ 94 cm, men; ≥ 80 cm, women), raised WHR (WHR: ≥ 1.0, men; ≥ 0.85, women), raised TG (TG ≥ 2.26 mmol/L), raised TC (TC ≥ 6.22 mmol/L), raised LDL (LDL ≥ 4.14 mmol/L), raised FPG (FPG≥ 7.0 mmol/L), raised SBP (SBP ≥ 140 mmHg), and raised DBP (DBP ≥ 90 mmHg).^18–21^

### SNP selection

We accessed all data (regardless of the ancestry of the population studied) for each trait in the NHGRI-EBI database of published genome-wide association studies (GWAS Catalog) as of May 25, 2019.^22^ The GWAS catalog is a curated comprehensive public repository of published GWAS reporting single nucleotide polymorphism (SNP)-trait associations with P-value <1 × 10^−5^. From the GWAS Catalog, we extracted the SNP identifier (RefSeq rs number) and the risk allele for each SNP reported. Each of the SNPs was then mapped to Ensembl release version 92 to identify the reference and alternate alleles. The set of overlapping SNPs between those extracted from the GWAS catalog and the target dataset were retained for constructing GRS. Further, we performed sensitivity analyses using independent SNPs obtained by pruning out the above SNPs with a variance inflation factor > 2 within a sliding “window” of size 50 bp shifted over 5 SNPs at every step.^23^

### Construction of GRS

An individual’s GRS was constructed as a weighted sum of the number of risk alleles over all the SNPs identified for each trait using PLINK 1.9.^24^ Effects sizes used for weighting were obtained from the UK Biobank^25^ for BMI, WC, HC, WHR, SBP, DBP, and T2D, or the largest study in the GWAS catalog for the other traits (Spracklen et al.^26^ for TC, TG, HDL, and LDL and Manning et al.^27^ for FPG). UK Biobank data were from White Bristish individuals, while Spracklen et al. study data were from European and East Asian indivuals. Manning et al. study data were from Europen-ancestry individuals. For FPG, GRS was constructed for non-T2D cases only. The sign of the effect size was appropriately flipped when the reported risk allele in the weight-source dataset was the alternate of the risk allele in the target dataset.

### Statistical analysis

Calibration of the GRS was assessed using correlations between GRS and traits and by plotting the observed mean or prevalence of a trait against its GRS deciles. Predictive utility of GRS was assessed using two metrics: (1) additional trait variability attributable to GRS in terms of adjusted R-Squared of the regression model, and, (2) additional discriminatory power attributable to GRS in terms of area under the Receiver Operating Characteristic Curve (AUC). R-squared assessments were based on comparisons of regression models fitted for each quantitative trait against traditional risk factors (age, sex, and principal components of ancestry and BMI [except when BMI was the trait under study]), with (GRS model) and without GRS (traditional model). Logistic regression models were fitted for T2D and Efron’s R^2^ used to estimate the additional variation in the probability of T2D explained by GRS.^28^ AUCs based on logistic regression models fitted for binary traits and additional discriminatory power of GRS were assessed by comparing the model of GRS plus traditional risk factors with the model of only traditional risk factors. All downstream analyses were performed in STATA version 15.1 (STATA Corp, Texas) and two-tailed value of P <0.05 were considered significant. The P-values referred to here relate to regresson and correlation coefficients of association between each trait and its corresponding GRS. These tests are not a ‘family of tests’ for which adjustment for a Family-wise Error Rate or other multiple testing adjustment is appropriate.^29^

## Results

### Distribution of GRS

Information about the cardiometabolic traits studied, number of SNPs, sources of weights and numbers of individuals studied are shown in **Table 1**. The number of SNPs used to construct GRS did not significantly differ between the three groups. The distribution of GRS for the cardiometabolic traits studied differed among the three groups, except for total cholesterol (TC) (**Figure 1**). Higher mean GRS were observed for type 2 diabetes (T2D) relative to other traits (Table 1), but this reflected a difference in weighting; the GRS for T2D was weighted using odds ratios while linear regression coefficients were used for the other traits. Overall, relative to AF and AA, EA had significantly higher GRS for six (waist circumference, WC; waist-hip ratio, WHR; systolic blood pressure, SBP; diastolic blood pressure, DBP; triglycerides, TG; low-density lipoprotein, LDL) out of the 12 traits studied. On the other hand, AF had a significantly higher GRS for hip circumference (HC), fasting plasma glucose (FPG) and T2D. The overlap of GRS distributions was greater between AF and AA (nearly identical for HC, WHR, TG, and high-density lipoprotein (HDL)) than between any one of them and EA, except for T2D and LDL for which there was greater overlap of GRS distributions between AF and EA. Generally, the GRS distributions among AA were consistently below or between the distributions among AF and EA.

**Table 1:**
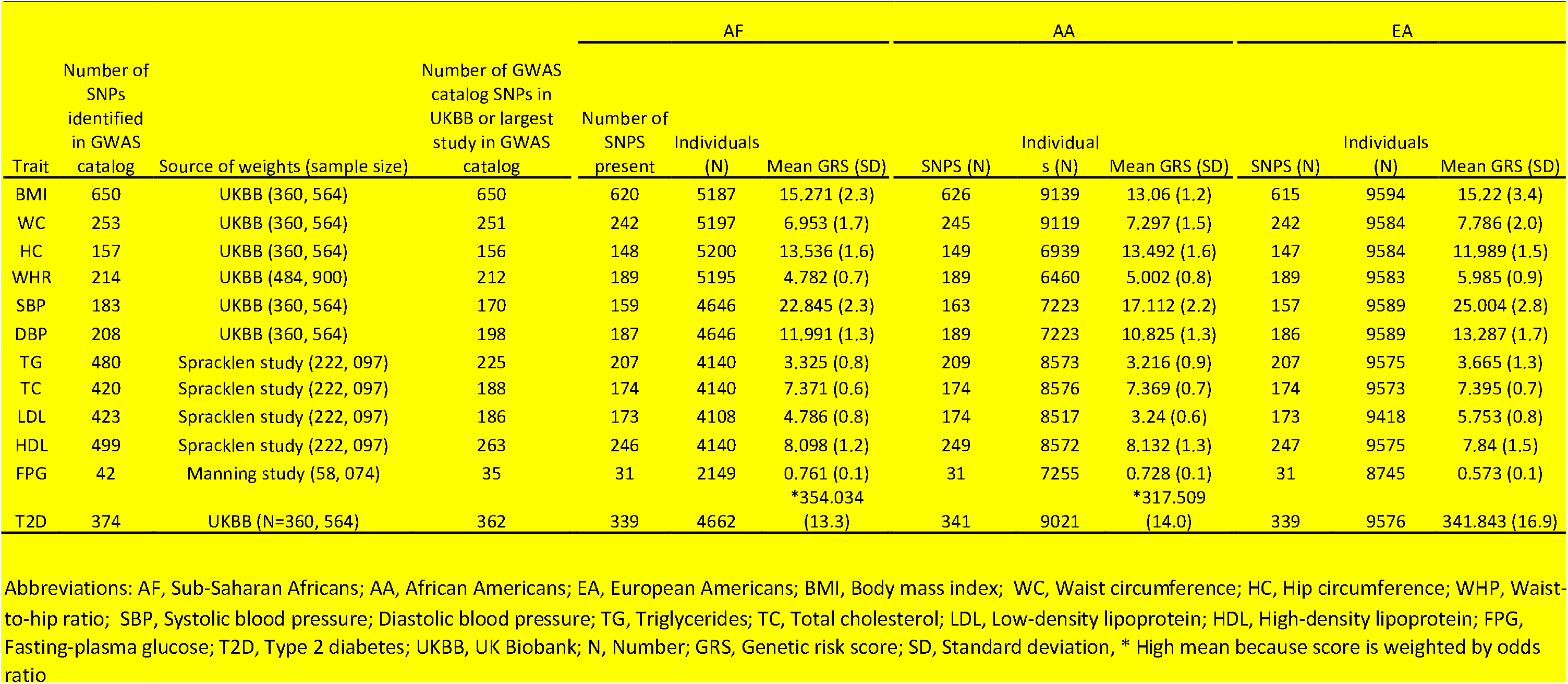
Sample size, descriptive summary of SNPs and source of weights.

**Figure 1:**
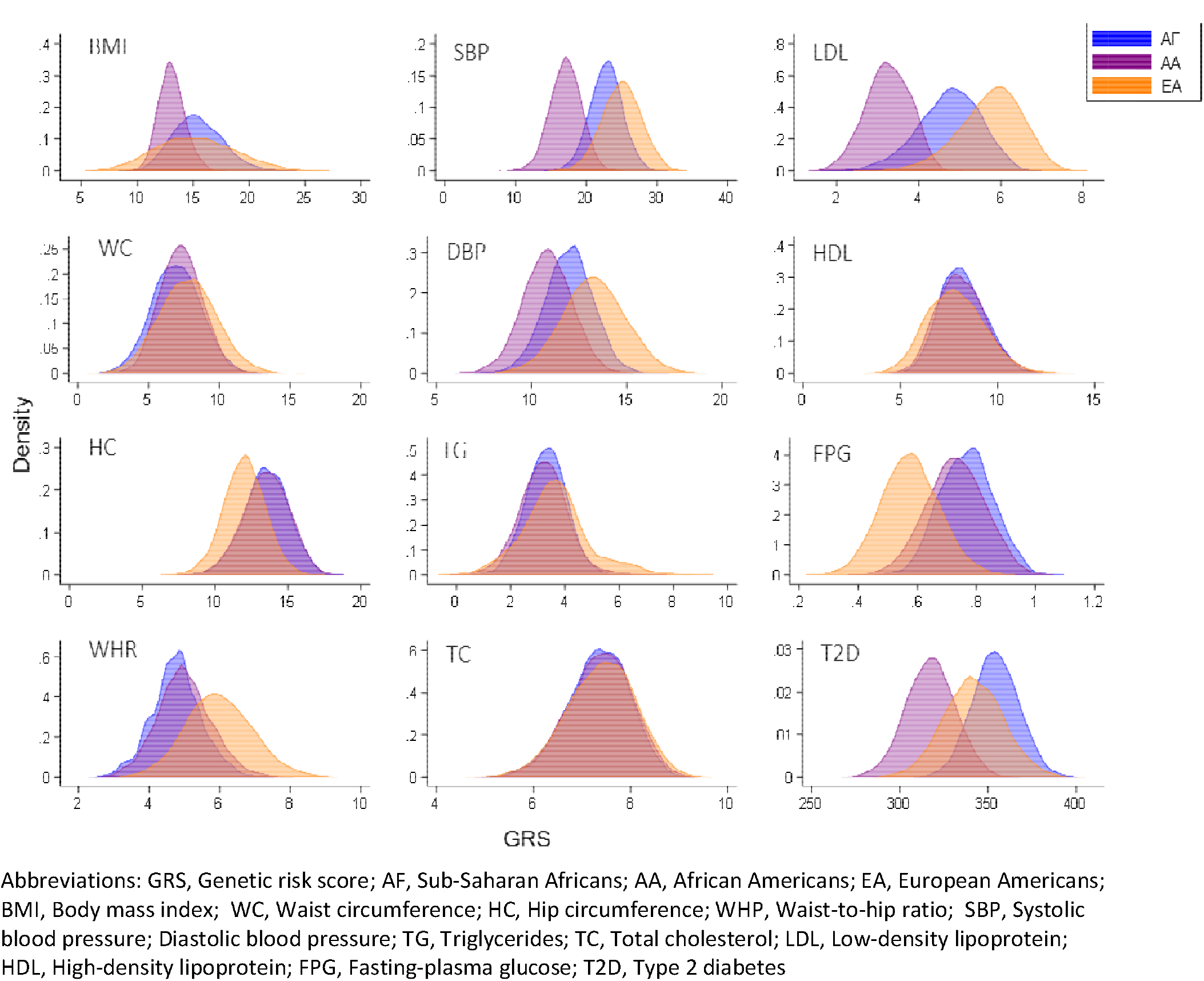
Distribution of genetic risk scores by group.

### Association of GRS with cognate outcomes

GRS were more strongly associated with their respective traits among EA relative to AF and AA (**Table 2**). Among EA, 11 of the 12 GRS-trait associations were statistically significant (P<0.001) while 10 and 8 of 12 GRS-trait associations were statistically significant among AA and AF, respectively (**Supplementary Figure 1**). In addition, the strongest GRS-trait associations were observed for lipid traits in all three groups.

**Table 2:** Correlations between traits and genetic risk score.

| Trait | AF |  | AA |  | EA |  |
| --- | --- | --- | --- | --- | --- | --- |
|  | Correlation coefficient | P-value | Correlation coefficient | P-value | Correlation coefficient | P-value |
| BMI | 0.051 | <0.001 | 0.041 | <0.001 | 0.091 | <0.001 |
| WC | 0.028 | 0.047 | 0.023 | 0.026 | 0.054 | <0.001 |
| HC | 0.043 | 0.002 | 0.033 | 0.006 | 0.081 | <0.001 |
| WHR | 0.012 | 0.376 | 0.007 | 0.564 | 0.010 | 0.342 |
| SBP | 0.008 | 0.583 | 0.034 | 0.004 | 0.071 | <0.001 |
| DBP | 0.023 | 0.122 | 0.049 | <0.001 | 0.062 | <0.001 |
| TG | 0.102 | <0.001 | 0.093 | <0.001 | 0.192 | <0.001 |
| TC | 0.113 | <0.001 | 0.124 | <0.001 | 0.186 | <0.001 |
| LDL | 0.141 | <0.001 | 0.095 | <0.001 | 0.186 | <0.001 |
| HDL | 0.100 | <0.001 | 0.199 | <0.001 | 0.182 | <0.001 |
| FPG | 0.001 | 0.962 | 0.020 | 0.061 | 0.084 | <0.001 |
| T2D | 0.111 | <0.001 | 0.069 | <0.001 | 0.100 | <0.001 |

Abbreviations: AF, sub-Saharan Africans; AA, African Americans; EA, European Americans; BMI, Body mass index; WC, Waist circumference; HC, Hip circumference; WHR, Waist-to-hip ratio; SBP, Systolic blood pressure; Diastolic blood pressure; TG, Triglycerides; TC, Total cholesterol; LDL, Low-density lipoprotein; HDL, High-density lipoprotein; FPG, Fasting-plasma glucose; T2D, Type 2 diabetes

### Predictive utility of GRS

In regression models adjusted for traditional risk factors and population genetic structure (represented by the first three principal components of ancestry), GRS was significantly associated with body mass index (BMI), DBP, lipid traits and T2D in all three groups (**Table 3**). Furthermore, among AA and EA, GRS was also significantly associated with WC, SBP and FPG, and, additionally, with HC among EA only. The effect sizes of the above seven GRS-trait associations (GRS association with BMI, DBP, lipid traits and T2D) ranked in roughly the same order, with the GRS-TC association being the strongest and GRS-BMI association the weakest. Notably, among these GRS-trait associations, the largest effect size was observed among EA for TG, TC, LDL and T2D, while the other three (BMI, DBP and T2D) had their largest effect sizes among AA. As an example, among GRS-trait associations common to all three groups, the GRS-TC association was the strongest and the effect sizes were 0.226, 0.216 and 0.281 mmol/l per unit increase in GRS (all P<0.0001) among AF, AA and EA, respectively. Furthermore, odds ratios for binary traits (comparing individuals in the top 10% of GRS with the rest) were statistically significant for lipids, FPG and T2D, except for raised TG and raised FPG among AF. (**Figure 2**).

**Table 3:** GRS effect size and R-squared.

**Panel A: AF**
| Trait | GRS Model |  |  |  | Model without GRS | Additional variation explained |
| --- | --- | --- | --- | --- | --- | --- |
|  | GRS effect size | SE | P value | R<sup>2</sup> | R<sup>2</sup> |  |
| BMI | 0.133 | 0.0336 | 0.0001 | 0.0767 | 0.0741 | 0.0026 |
| WC | 0.074 | 0.0676 | 0.2749 | 0.5700 | 0.5700 | 0.0000 |
| HC | 0.095 | 0.0722 | 0.1898 | 0.5545 | 0.5545 | 0.0000 |
| WHR | 0.0004 | 0.0015 | 0.7781 | 0.1965 | 0.1967 | -0.0002 |
| SBP | 0.141 | 0.1383 | 0.3068 | 0.1640 | 0.1640 | 0.0000 |
| DBP | 0.349 | 0.1514 | 0.0213 | 0.0659 | 0.0651 | 0.0008 |
| TG | 0.073 | 0.016 | <0.0001 | 0.1803 | 0.1761 | 0.0042 |
| TC | 0.272 | 0.036 | <0.0001 | 0.0628 | 0.0502 | 0.0126 |
| LDL | 0.226 | 0.025 | <0.0001 | 0.0781 | 0.0596 | 0.0185 |
| HDL | 0.041 | 0.006 | <0.0001 | 0.0403 | 0.0293 | 0.0110 |
| FPG | -0.1701 | 0.1570 | 0.2788 | 0.0447 | 0.0447 | 0.0000 |
| T2D | 1.020 | 0.0024 | <0.0001 | 0.1180 | 0.1050 | 0.0130 |

**Panel B: AA**
| Trait | GRS Model |  |  |  | Model without GRS | Additional variation explained |
| --- | --- | --- | --- | --- | --- | --- |
|  | GRS effect size | SE | P-value | R<sup>2</sup> | R<sup>2</sup> |  |
| BMI | 0.242 | 0.0641 | 0.0002 | 0.0214 | 0.0200 | 0.0014 |
| WC | 0.122 | 0.0558 | 0.0284 | 0.7679 | 0.7678 | 0.0001 |
| HC | -0.014 | 0.0437 | 0.7436 | 0.8387 | 0.8387 | 0.000 |
| WHR | 0 | 0.0012 | 0.7457 | 0.2430 | 0.2431 | -0.0001 |
| SBP | 0.334 | 0.0997 | 0.0008 | 0.1036 | 0.1023 | 0.0013 |
| DBP | 0.448 | 0.1041 | <0.0001 | 0.0174 | 0.0150 | 0.0024 |
| TG | 0.089 | 0.0101 | <0.0001 | 0.0359 | 0.0272 | 0.0087 |
| TC | 0.216 | 0.0181 | <0.0001 | 0.0651 | 0.0496 | 0.0155 |
| LDL | 0.187 | 0.02 | <0.0001 | 0.0462 | 0.0365 | 0.0097 |
| HDL | 0.066 | 0.0034 | <0.0001 | 0.0980 | 0.0591 | 0.0389 |
| FPG | 0.261 | 0.0724 | 0.0003 | 0.1806 | 0.1793 | 0.0013 |
| T2D | 1.014 | 0.0022 | <0.0001 | 0.0670 | 0.0620 | 0.0050 |

**Panel C: EA**
| Trait | GRS Model |  |  |  | Model without GRS | Additional variation explained |
| --- | --- | --- | --- | --- | --- | --- |
|  | GRS effect size | SE | P-value | R<sup>2</sup> | R<sup>2</sup> |  |
| BMI | 0.124 | 0.0137 | <0.0001 | 0.0153 | 0.0070 | 0.0083 |
| WC | 0.077 | 0.0282 | 0.0066 | 0.8215 | 0.8214 | 0.0001 |
| HC | 0.104 | 0.0295 | 0.0004 | 0.8027 | 0.8025 | 0.0002 |
| WHR | 0 | 0.0006 | 0.71 | 0.4750 | 0.4750 | 0.00 |
| SBP | 0.439 | 0.0569 | <0.0001 | 0.1438 | 0.1385 | 0.0053 |
| DBP | 0.369 | 0.059 | <0.0001 | 0.0896 | 0.0860 | 0.0036 |
| TG | 0.15 | 0.0077 | <0.0001 | 0.1085 | 0.0737 | 0.0348 |
| TC | 0.281 | 0.0147 | <0.0001 | 0.0721 | 0.0369 | 0.0352 |
| LDL | 0.237 | 0.0125 | <0.0001 | 0.0636 | 0.0280 | 0.0356 |
| HDL | 0.05 | 0.0025 | <0.0001 | 0.3061 | 0.2767 | 0.0294 |
| FPG | 0.707 | 0.0509 | <0.0001 | 0.1374 | 0.1184 | 0.0190 |
| T2D | 1.023 | 0.0023 | <0.0001 | 0.0690 | 0.0550 | 0.0140 |

AF, sub-Saharan Africans; AA, African Americans; EA, European Americans; GRS effect size= $\beta$ Linear regression coefficient (for quantitative trait) or odds ratio (for disease trait); SE, Standard error; R<sup>2</sup>, Adjusted R-squared; GRS Model: Trait ∝ age+sex+BMI+PC1+PC2+PC3+GRS (BMI excluded in covariates when it's the trait under study); BMI, Body mass index; WC, Waist circumference; HC, Hip circumference; WHR, Waist-to-hip ratio; SBP, Systolic blood pressure; DBP, Diastolic blood pressure; TG, Triglycerides; TC, Total cholesterol; LDL, Low-density lipoprotein; HDL, High-density lipoprotein; FPG, Fasting-plasma glucose; T2D, Type 2 diabetes; PC1, PC2, PC3, the first three principal components of ancestry; Additional variation explained is in percentage points

**Figure 2:**
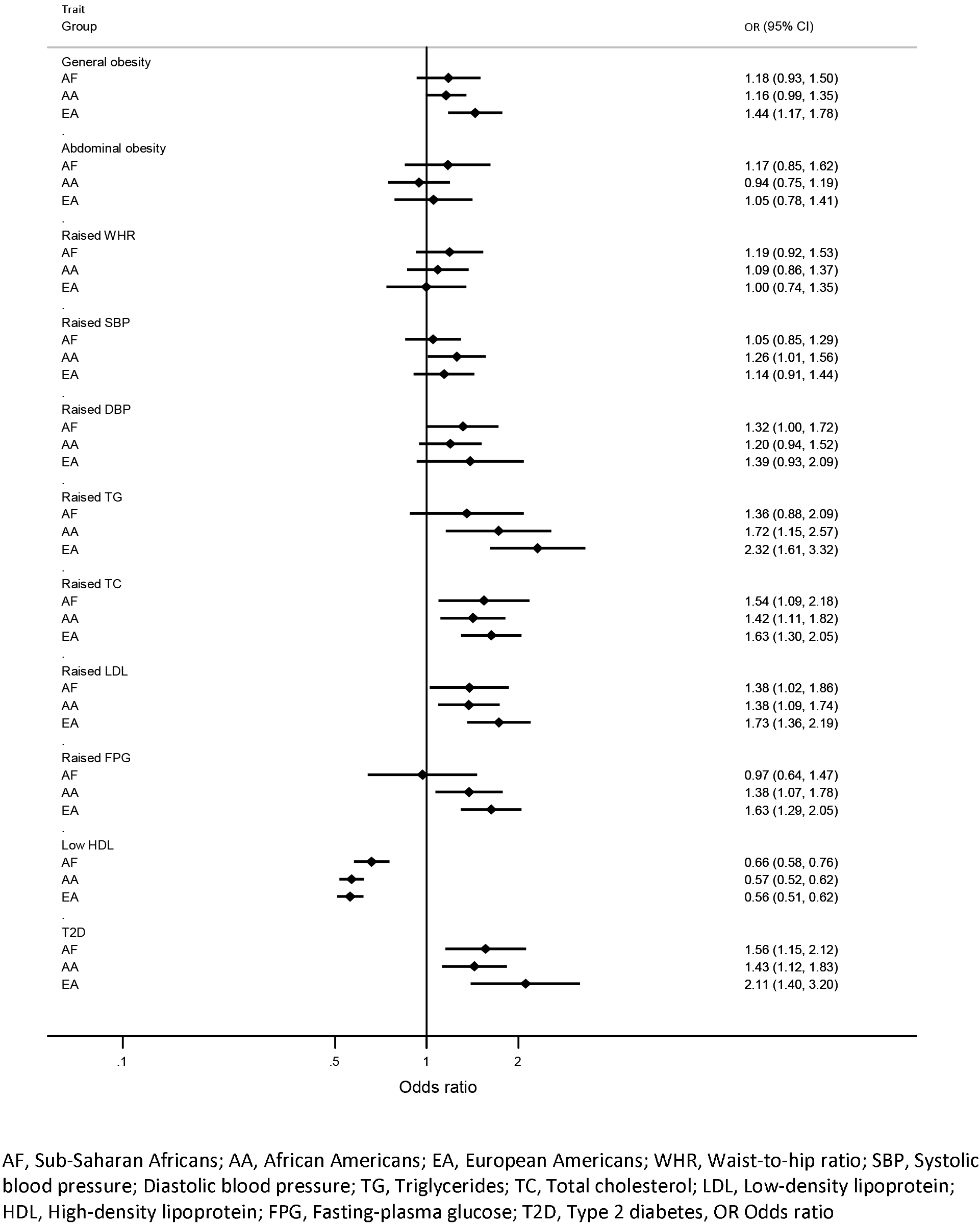
Association between GRS (individuals in the top 10% versus the others) and binary traits.

The predictive utility of GRS was assessed in terms of additional variation explained by the model including GRS (GRS model) relative to variation explained by the model of traditional risk factors only (traditional model). The predictive utility of GRS showed significant variation both among traits and among groups (**Figure 3**). We observed substantial predictive utility of GRS for lipid traits and T2D in all groups, and additionally among EA for BMI and FPG. Among AA, GRS also appeared to have predictive power for DBP. However, the predictive power of GRS was significantly greater in EA compared with AF and AA, showing up to 5-fold and 20-fold greater predictive utility of GRS in EA relative to AA and AF, respectively. However, exceptions were observed for HDL and DBP, for which the predictive utility of GRS was greater among AF (HDL, 4-fold) and AA (HDL, 6-fold; DBP, 3.8-fold) compared with EA. Between AF and AA, disparity in the predictive value of GRS was less consistent and less profound, but still substantial for some traits. For example, the predictive utility of GRS for TG was 13-fold greater among AA relative to AF but 1.5-fold greater among AF relative to AA for HDL.

**Figure 3:**
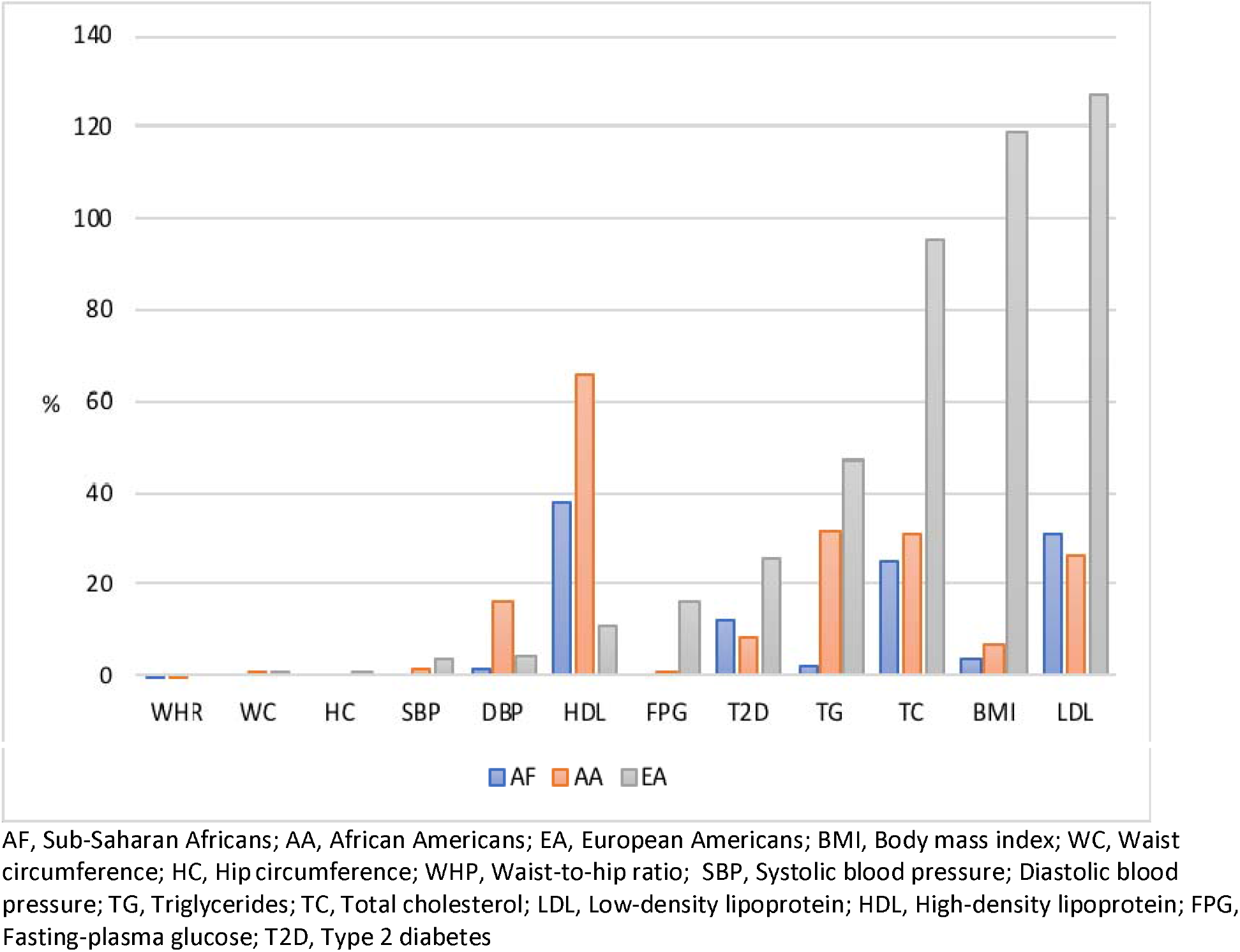
Percentage increase in R-squared attributable to genetic risk score.

The predictive utility of GRS based on additional trait variation explained was limited for traits whose variability was substantially explained by the model of traditional risk factors. This was especially true for anthropometric traits across groups except for BMI among EA. Whereas GRS increased the proportion of variation explained in BMI by only 3.5% and 7.0% among AF and AA, respectively, the corresponding increase was 118.6% among EA, representing a 34-fold and 17-fold greater predictive utility in EA compared with AF and AA, respectively.

We assessed the predictive utility of GRS for dichotomized transformations of the quantitative traits in addition to T2D using Area Under the Receiver Operating Characteristic curve (AUC). The heterogeneity among traits and disparity among groups of the predictive utility of GRS were similar under this approach. We observed a substantial predictive utility of GRS for components of lipid dysregulation and T2D across groups but more so among EA (**Figure 4**). Among AF, the greatest increases in AUC (lesss than 2% gains) were observed for lipid traits and T2D. Among AA, lipid traits and T2D had increases of up to 5.7%. Among EA, increases up to 23.2% were observed for nine of 12 traits, again showing predictive disparities in favor of EA.

**Figure 4:**
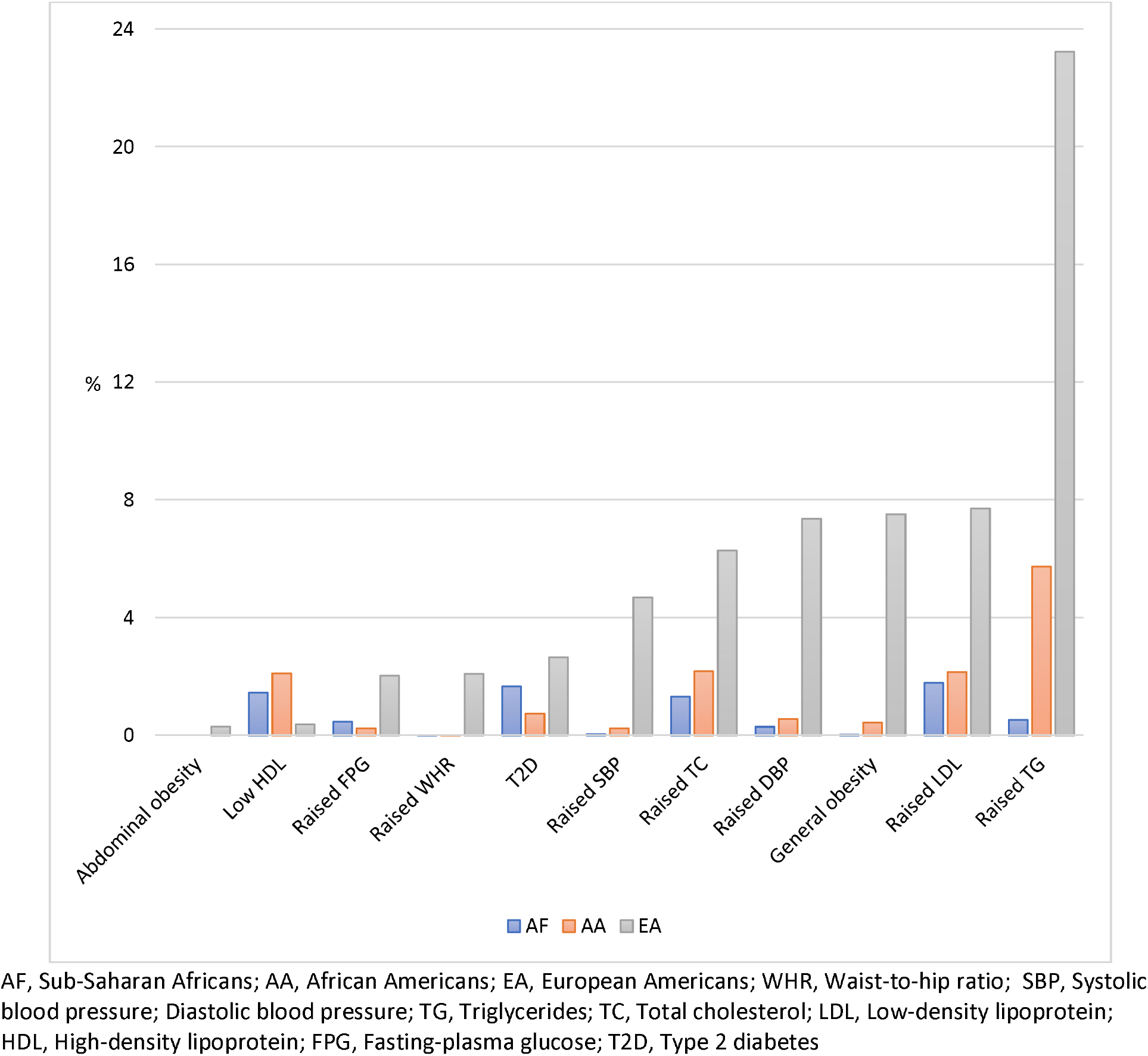
Percentage increase in Area Under the Receiver Operating Characteristic curve (AUC) attributable to genetic risk score.

These figures correspond to a greater predictive utility of GRS among EA compared with AF of 116-, 44-, 25-, 5-, 4-, and 4-fold, for raised SBP, raised DBP, raised TC, raised LDL, raised TG and raised FPG, respectively. In the single instance where the disparity was in favor of AF relative to EA, the predictive utility of GRS for low HDL was 4-fold greater among AF compared with EA. Comparing EA with AA, the greatest disparities in favor of EA were observed for SBP (20-fold), raised FPG (8-fold), raised TG, raised LDL and raised T2D (4-fold). Between AF and AA, the greatest disparities were observed in favor of AA as follows: raised TG (11-fold), raised SBP (6-fold) and low HDL (1.4-fold); while disparities in favor of AF relative to AA were relatively modest, in that the predictive utility of GRS was 2-fold greater for raised FPG and raised T2D among AF.

As with the R-squared method, we found limited predictive utility of GRS among AF and AA under the AUC approach for traits such as general obesity and abdominal obesity, whether defined by WC or WHR. Thus, disparity in predictive utility of GRS in favor of EA relative to AF were extremely large for these traits. For example, the predictive utility of GRS for general obesity and raised WHR was 249- and 172-fold, respectively, greater among EA compared with AF. The disparity was reduced between EA and AA for general obesity but not for raised WHR, thus the predictive utility of GRS was 17- and 172-fold, respectively, greater among EA compared with AA. For abdominal obesity, where GRS had no predictive utility beyond traditional risk factors among AF and AA, the predictive disparity in favor of EA was infinite.

### Sensitivity analyses

As a sensitivity analysis, we assessed the predictive utility of GRS constructed from only independent SNPs (*i.e.*, with SNPs in high linkage disequilibrium (LD) removed) (*pruned*GRS). The predictive utility of *pruned*GRS broadly recapitulated the above results: consistent GRS-trait associations for lipids with greater predictive power among EA compared with AF and AA (**Supplementary Figure 2**). Predictive utility was lower for *pruned*GRS compared with GRS based on all SNPs in all three groups except for LDL among AF and AA. The number of SNPs removed due to high LD was lower for AF compared with AA and EA across traits, but largely comparable between AA and EA (**Supplementary Table 1**).

## Discussion

Using a dataset of about 24,000 individuals, we demonstrate that the predictive utility of GRS showed significant variation among 12 cardiometabolic traits and among populations with differing proportion of African ancestry and in comparison with European ancestry populations. GRS was well calibrated for lipids in all three groups but was only calibrated well for the other traits in EA. Additionally, the predictive utility of GRS was often strongest in EA and poorest among AF. Between AF and AA, differences in GRS performance were less pronounced but tended to favor AA, perhaps reflecting European admixture in AA. To our knowledge, this is the first study of GRS for complex traits in continental Africans and the first comparison of GRS predictive utility between continental Africans and African Americans. These findings have important implications for the potential benefits to be derived from the application of GRS in routine clinical risk prediction across populations of different ancestries.

The variation in the predictive performance of GRS among traits likely reflects differential heritability—a measure of the relative influence of genetic and environmental factors on a trait. The predictive power of GRS has been shown to correlate with heritability and greater heritability has been reported for lipids compared with obesity/anthropometric, blood pressure and glycemic traits.^30,31^ This is consistent with the observations from the present study in which lipid traits stood out in terms of association with GRS. However, we note that among EA, the predictive utility of GRS was higher for BMI than some lipid components, suggesting that differences in heritability among traits may not be consistent across populations due to varying gene-environmental interactions. Additionally, among EA, GRS showed no predictive utility for WHR and WC under the additional explained phenotypic-variation approach but showed some utility, although limited, under the discriminatory power approach. This observation is likely the consequence of minimal variability in quantitative WHR versus the substantial variance of the binomial distribution of the corresponding binary transformation.

The predictive utility of GRS among AA was better than in AF but worse than in EA in the present study. Reduced prediction accuracy in AA relative to EA is consistent with previous reports of lower predictive utility of similarly constructed GRS in admixed individuals compared with Europeans.^5,7,9,32^ The observed pattern of predictive performance of GRS is consistent with the disproportionately large number of individuals of European ancestry in current genome-wide discovery studies and the degree of genetic divergence of AF and AA from EA. EA contribute nearly four-fifths of individuals included in current GWAS, and AF is more genetically distant from EA than admixed AA, who have about 20% European ancestry.^33^ In addition, under-representation of diverse global populations in available genomic resources (including genotyping arrays and imputation panels) means that these resources do not adequately capture global genetic diversity due to differences in allele frequencies and linkage disequilibrium patterns among populations.^9,32,34,35^ When population differences in variant effect sizes are factored in, an expected consequence is poorer prediction accuracy of GRS in the underrepresented populations. These considerations highlight the need for genomic resources, methods, and tools that take into account global genetic diversity. Indeed, there is increasing evidence demonstrating improved GRS predictive accuracy when GRS are constructed from ancestry-matched variants and GWAS summary statistics.^9,36,37^ Other factors that are important in disparities of GRS predictive utility include differences in polygenic adaptation due to natural selection, historical population size, residual uncorrected population structure and etiological differences between populations.^38–41^ Other possible factors include differences in genetic architecture due to gene-environment or gene-gene interactions in admixed populations or monomorphism of the causal variant in an ancestral population.^42,43^ In this regard, it is important to note that AF differ from AA not just in genetic variation but also in environmental factors that influence cardiometabolic phenotypes, including dietary, behavioral, socio-economic, and other lifestyle factors.^44^

The intriguing lack of predictive utility of GRS for TG among AF is unclear but parallels the existence of lower TG observed in African ancestry individuals compared with non-African-ancestry individuals.^45^ A significant role for a genetic influence characterized by ancestry-specific loci has been suggested because of the consistency of lower TG levels across African-ancestry populations despite divergent environmental contexts and the persistence of lower TG among AA compared with EA in spite of similar environments.^33,46^ Therefore, poor predictive utility of GRS for TG among AF may be a reflection of non-transferability of current GWAS loci to AF possibly due to differences in sample size, effect size, allele frequency and gene-environment interactions. For HDL, the role of a genetic influence is less clear because of inconsistent differences in HDL levels among populations of different ancestry. Whereas AA tend to have higher HDL levels compared with EA, AF from West Africa have been shown to have lower levels of HDL, suggesting an important role for environmental factors.^33^ Further research is needed to clarify the potential role of underlying genetic differences as the force behind HDL variation among populations of different ancestry and its impact on the predictive utility of GRS in the context of environmental differences.

Inspite of concerns about the impact on health disparities, our findings are indicative of a promising role for GRS in predicting the risk of hypercholesteroleamia across populations of different ancestral backgrounds. A potential application of GRS in this context could be assessing additive risk of elevated LDL beyond the causative monogenic mutations of Familial Hypercholesteroleamia (FH). As high GRS has been shown to be associated with severity of the FH phenotype, carriers of monogenic FH-mutations with etreme GRS could be prioritsied for early intervention including treatment with statins, while knowledge of concomitant high GRS could encourage adherence to treatment among FH patients.^47,48^

Important strengths of this study are the large sample size, use of independent datasets for discovery, and assessment of predictive utility in different populations. Additionally, SNPs were identified from the NHGRI-EBI GWAS Catalog (a curated comprehensive public repository of published GWAS) and highly precise summary statistics used for weighting were obtained from the UK Biobank, which has genotype and extensive phenotypic data on ~500,000 individuals.^22,25^ However, our findings should be interpreted in the context of the limitations of the study. First, we only included SNPs that reached the GWAS-Catalog criteria of 1 × 10^−5^ level of significance for constructing GRS. It is possible that there are SNPs not yet identified with the current sample sizes, but which may be associated with the traits studied. Second, we did not account for gene-gene and gene-environment interactions, which may limit the predictive utility of GRS. Finally, the predictive utility of GRS observed in this study might be understated if most of the variants used to construct GRS do not tag the causal variants for the traits studied. This is particularly relevant because LD is weaker in African-ancestry individuals compared with European-ancestry individuals in whom the majority of current genetic variants were discovered.

This first evaluation of GRS in continental Africans demonstrates that the predictive performance of GRS for cardio-metabolic traits is markedly poor among sub-Saharan Africans and currently provides little or no benefit over traditional risk factors. We also confirm that GRS is worse in African Americans compared to European Americans. Therefore, unlike for EA populations, GRS for cardiometabolic disorders remain suboptimal for clinical translation in continental Africans as well as in African Americans. These findings add to the growing understanding of the strengths and limitations of the applications of GRS in routine clinical and/or public health settings and highlights the need to increase the inclusion of underrepresented populations in genomic discovery to promote equity in translation of such discovery.

## Acknowledgements/Funding

The contents of this publication are solely the responsibility of the authors and do not necessarily represent the official view of the National Institutes of Health (NIH). This research was supported by the Intramural Research Program of the Center for Research on Genomics and Global Health (CRGGH). The CRGGH is supported by the National Human Genome Research Institute, the National Institute of Diabetes and Digestive and Kidney Diseases and the Office of the Director at the National Institutes of Health (1ZIAHG200362). This work used the computational resources of the NIH HPC Biowulf cluster (https://hpc.nih.gov).

The following studies were funded by the listed NIH grants.

AADM was supported by NIH grant 3T37TW00041-03S2.

HUFS was supported by NIH grants S06GM008016-320107 (Dr. Rotimi), S06GM008016-380111 (Dr. Adeyemo), and 2M01RR010284. This research was also supported in part by the NIH Intramural Research Program in the Center for Research on Genomics and Global Health (1ZIAHG200362).

ARIC was supported by NIH grants N01-HC-55015, N01-HC-55018, N01-HC-55016, N01-HC-55021, N01-HC-55019, N01-HC-55020, and N01-HC-55022.

CFS was supported by NIH grants R01-HL-46380 and M01-RR-00080.

JHS was supported by NIH grants N01-HC-95170, N01-HC-95171, and N01-HC-95172.

MESA was supported by NIH grants N01-HC-95159, N01-HC-95160, N01-HC-95161, N01-HC-95162, N01-HC-95168, N01-HC-95163, N01-HC-95164, N01-HC-95165, N01-HC-95166, N01-HC-95167, N01-HC-95169, and R01-HL-071205.

The funders had no role in study design, data collection, data analysis, interpretation, or writing of the paper.

## Authors’ contributions

CR, AA, conceptualized the study; KE, GC, JZ, and DS performed data management and statistical analysis; KE and AA drafted the paper; CR, AA, and KE edited the paper. All contributors reviewed and approved the manuscript.

