## Supplemental Tables and Figures for "Genetic Risk Scores for Cardiometabolic Traits in Sub-Saharan African Populations"

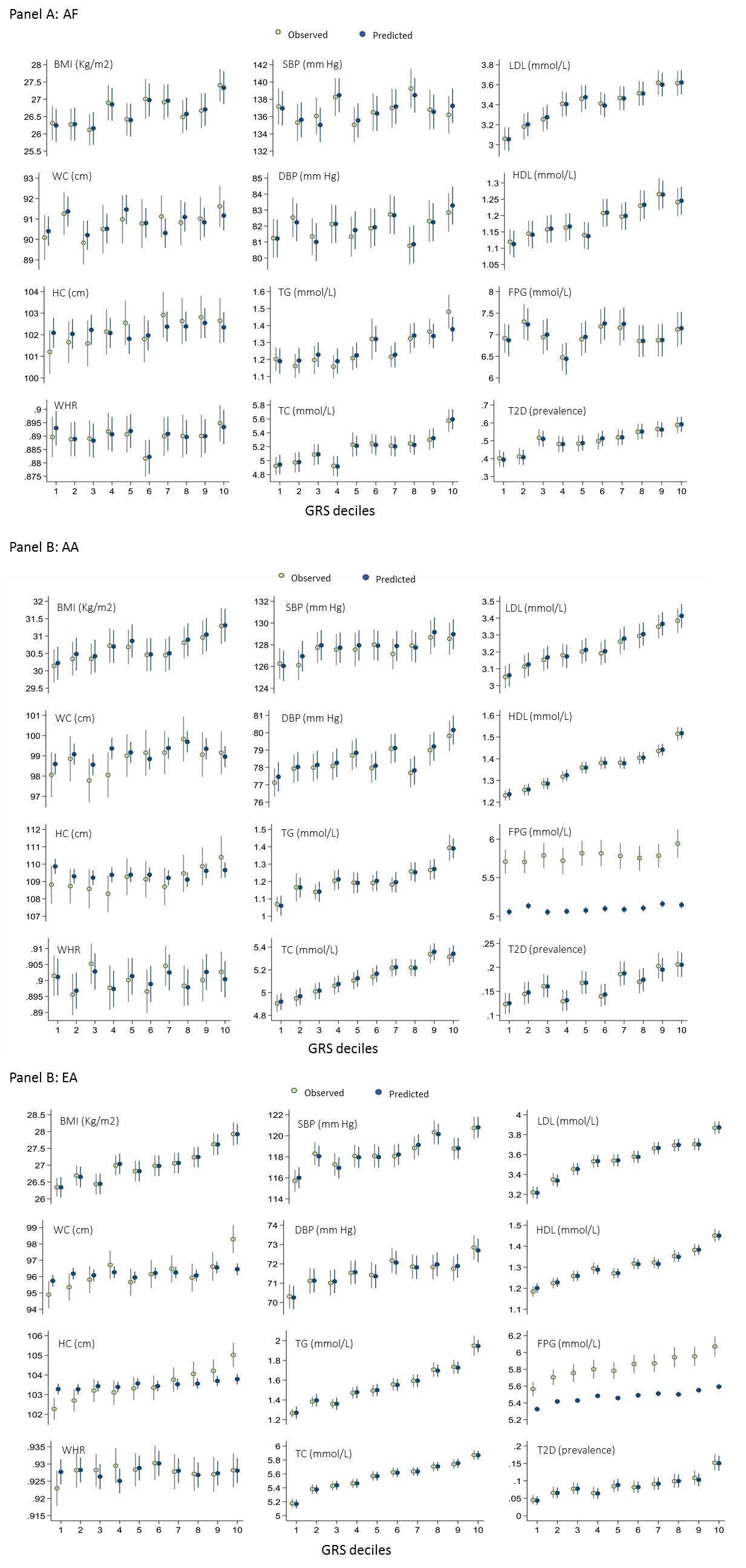

GRS, Genetic risk score; AF, Sub-Saharan Africans; AA, African Americans; EA, European Americans; BMI, Body mass index; WC, Waist circumference; HC, Hip circumference; WHP, Waist-to-hip ratio; SBP, Systolic blood pressure; Diastolic blood pressure; TG, Triglycerides; TC, Total cholesterol; LDL, Low-density lipoprotein; HDL, High-density lipoprotein; FPG, Fasting-plasma glucose; T2D, Type 2 diabetes; Error bars correspond to 95% CI

Supplementary Figure 1: Observed and predicted mean (prevalence for binary trait) across genetic risk score deciles.

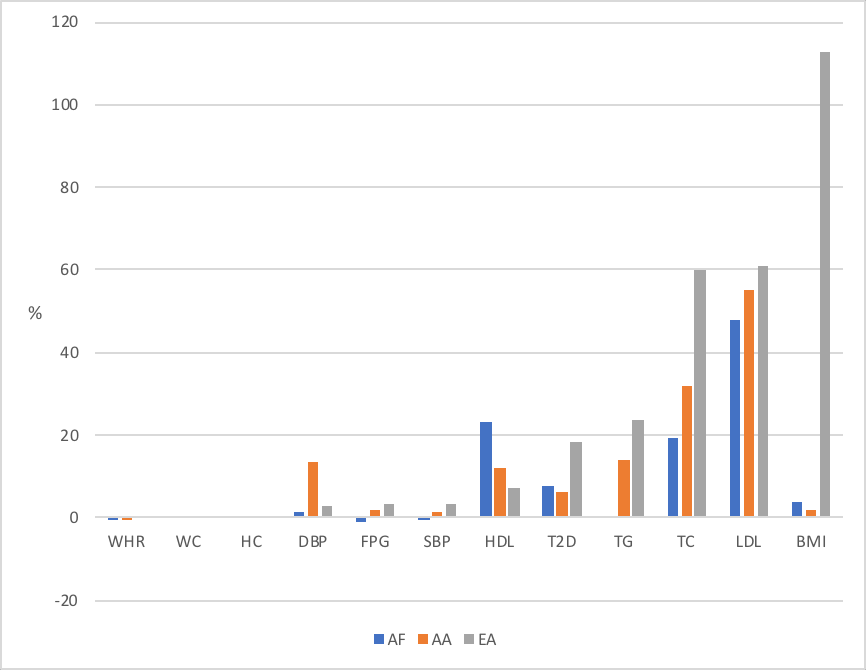

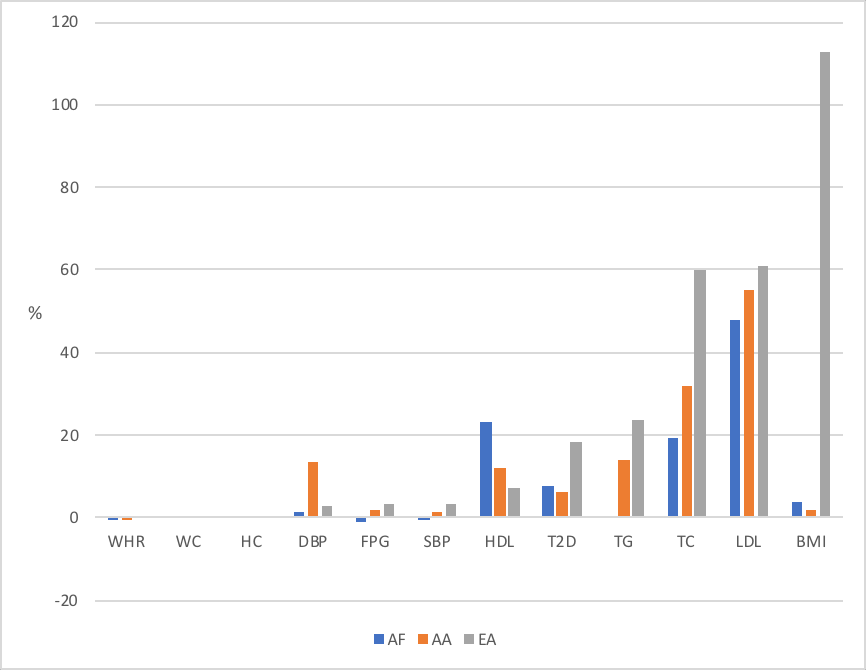

AF, Sub-Saharan Africans; AA, African Americans; EA, European Americans; BMI, Body mass index; WC, Waist circumference; HC, Hip circumference; WHP, Waist-to-hip ratio; SBP, Systolic blood pressure; Diastolic blood pressure; TG, Triglycerides; TC, Total cholesterol; LDL, Low-density lipoprotein; HDL, High-density lipoprotein; FPG, Fasting-plasma glucose; T2D, Type 2 diabetes

Supplementary Figure 2: Percentage increase in R-squared attributable to genetic risk score constructed from approximately independent SNPs (*prunedGRS*).

Supplementary Table 1: Number and descriptive summary statistics of independent SNPs remaining after pruning.

|  |  | AF | | |  | AA | | |  | EA | | |
| --- | --- | --- | --- | --- | --- | --- | --- | --- | --- | --- | --- | --- |
| Trait |  | SNPS (N) | Individuals (N) | Mean GRS (SD) |  | SNPS (N) | Individuals (N) | Mean GRS (SD) |  | SNPS (N) | Individuals (N) | Mean GRS (SD) |
| BMI |  | 407 | 5187 | 15.271 (2.3) |  | 365 | 9139 | 13.06 (1.2) |  | 377 | 9594 | 15.22 (3.4) |
| WC |  | 167 | 5197 | 6.953 (1.7) |  | 154 | 9119 | 7.297 (1.5) |  | 157 | 9584 | 7.786 (2.0) |
| HC |  | 135 | 5200 | 13.536 (1.6) |  | 121 | 6939 | 13.492 (1.6) |  | 123 | 9584 | 11.989 (1.5) |
| WHR |  | 124 | 5195 | 4.782 (0.7) |  | 119 | 6460 | 5.002 (0.8) |  | 113 | 9583 | 5.985 (0.9) |
| SBP |  | 136 | 4646 | 22.845 (2.3) |  | 123 | 7223 | 17.112 (2.2) |  | 125 | 9589 | 25.004 (2.8) |
| DBP |  | 159 | 4646 | 11.991 (1.3) |  | 148 | 7223 | 10.825 (1.3) |  | 149 | 9589 | 13.287 (1.7) |
| TG |  | 130 | 4140 | 3.325 (0.8) |  | 120 | 8573 | 3.216 (0.9) |  | 119 | 9575 | 3.665 (1.3) |
| TC |  | 113 | 4140 | 7.371 (0.6) |  | 110 | 8576 | 7.369 (0.7) |  | 101 | 9573 | 7.395 (0.7) |
| LDL |  | 107 | 4108 | 4.786 (0.8) |  | 97 | 8517 | 3.24 (0.6) |  | 95 | 9418 | 5.753 (0.8) |
| HDL |  | 139 | 4140 | 8.098 (1.2) |  | 127 | 8572 | 8.132 (1.3) |  | 123 | 9575 | 7.84 (1.5) |
| FPG |  | 27 | 2149 | 0.761 (0.1) |  | 21 | 7255 | 0.728 (0.1) |  | 23 | 8745 | 0.573 (0.1) |
| T2D |  | 211 | 4662 | 354.034 (13.3) |  | 188 | 9021 | 317.509 (14.0) |  | 192 | 9576 | 341.843 (16.9) |

Abbreviations: AF, Sub-Saharan Africans; AA, African Americans; EA, European Americans; BMI, Body mass index; WC, Waist circumference; HC, Hip circumference; WHP, Waist-to-hip ratio; SBP, Systolic blood pressure; Diastolic blood pressure; TG, Triglycerides; TC, Total cholesterol; LDL, Low-density lipoprotein; HDL, High-density lipoprotein; FPG, Fasting-plasma glucose; T2D, Type 2 diabetes; N, Number; GRS, Genetic risk score; SD, Standard deviation
